# Atad3 is Essential for Mitochondrial Permeability Transition Pore Opening and Cardiac Ischemia Reperfusion Injury

**DOI:** 10.1101/2025.06.13.658955

**Authors:** Dexter Robichaux, Sual J Lopez, Arielys Mendoza, Daniel Ramirez, Suh-Chin J Lin, Jeffery D Molkentin, Pablo M Peixoto, Jason Karch

## Abstract

The molecular identity of the mitochondrial permeability transition pore (mPTP) has remained elusive since the discovery of its existence over 75 years ago. Despite the numerous candidate proteins proposed, none have withstood genetic ablation, leaving them relegated to auxiliary regulatory roles. To date, no essential mPTP component has been identified. Here, we establish ATAD3 as the first essential component of the mPTP. Genetic deletion of Atad3 in cardiomyocytes and hepatocytes renders heart and liver mitochondria incapable of undergoing Ca^2+^-induced mPTP-dependent swelling. Moreover, these mitochondria exhibit the highest Ca^2+^ retention capacity ever reported following genetic perturbation of the mPTP. Furthermore, patch-clamp recordings of recombinant ATAD3a in liposomes reveal intrinsic channel activity. Given the established role of mPTP-dependent necrosis in driving ischemia/reperfusion (I/R) injury, we show that cardiac-specific Atad3 deletion markedly reduces infarct size following I/R, with no additive protection from cyclosporine A. Together, these findings establish ATAD3 as an core, putative pore-forming component essential for mPTP opening and mPTP-dependent necrosis, resolving a long-standing mystery in mitochondrial biology.

## Introduction

The mitochondrial permeability transition pore (mPTP) is a Ca^2+^-activated channel in the inner mitochondrial membrane that leads to mitochondrial depolarization, organelle swelling, and necrotic cell death^1,2^. Despite decades of research, the molecular identity of the pore remains elusive^3^. While several proteins, including cyclophilin D (CypD), adenine nucleotide translocators (ANTs), and Bax/Bak1, have been implicated, genetic deletion of each fails to completely abrogate pore activity^4-10^. However, deletion of each of these regulators confer mPTP desensitization and protection from myocardial ischemia/reperfusion (I/R) injury ^4,5,9-11^. Consequently, these factors are regarded as regulators, rather than essential components.

ATAD3 is a mitochondrial AAA+ ATPase is an integral inner membrane protein that spans the inner membrane space and contains a substantial matrix domain^12^. Originally, the mPTP was described as an inner membrane pore that spans the inner membrane space^13,14^. Although the function of ATAD3 is under current investigation, it has been previously linked to mitochondrial nucleoid organization and cristae structure and its oligomerization has been associated with nureodegeneration^15-17^. We hypothesize that ATAD3 may contribute to mPTP formation based on its location, topology, and potential ability to oligmerize within the inner mitochondrial membrane. Here, we demonstrate that ATAD3 is essential for mPTP activity in both cardiac and hepatic mitochondria, and its loss confers profound resistance to ischemia/reperfusion (I/R) injury in vivo.

## Results

In order to determine the function of ATAD3, we employed a genetic loss of function approach to investigate how the loss of Atad3 effects mitochondrial and mPTP function as well as I/R injury. Here, we generated mice deficient for ATAD3 in hepatocytes or cardiomyocytes using a CRE LOXP strategy (Fig. 1A and B). Tissue panel analysis revealed that ATAD3 is ubiquitously expressed in mice (Fig. 1C). To confirm deletion of Atad3, we performed western blot analysis on isolated mitochondria from both livers and hearts isolated from Atad3 tissue specific deleted mice and controls (Fig. 1D). At 3 months of age, deletion of ATAD3 from the hearts did not result in changes in cardiac function or size assessed by histology, gravimetric analyses, and echocardiography (Fig. 1E-G). Notably, at 8 months of age, the Atad3 deleted hearts have increase in size, reduced function, and increased interstitial fibrosis (Fig. H-J). Furthermore, cardiac specific deletion of Atad3 leads to shorten life expectancy as these mice die by 9 months of age (Fig. K).

**Fig. 1.**
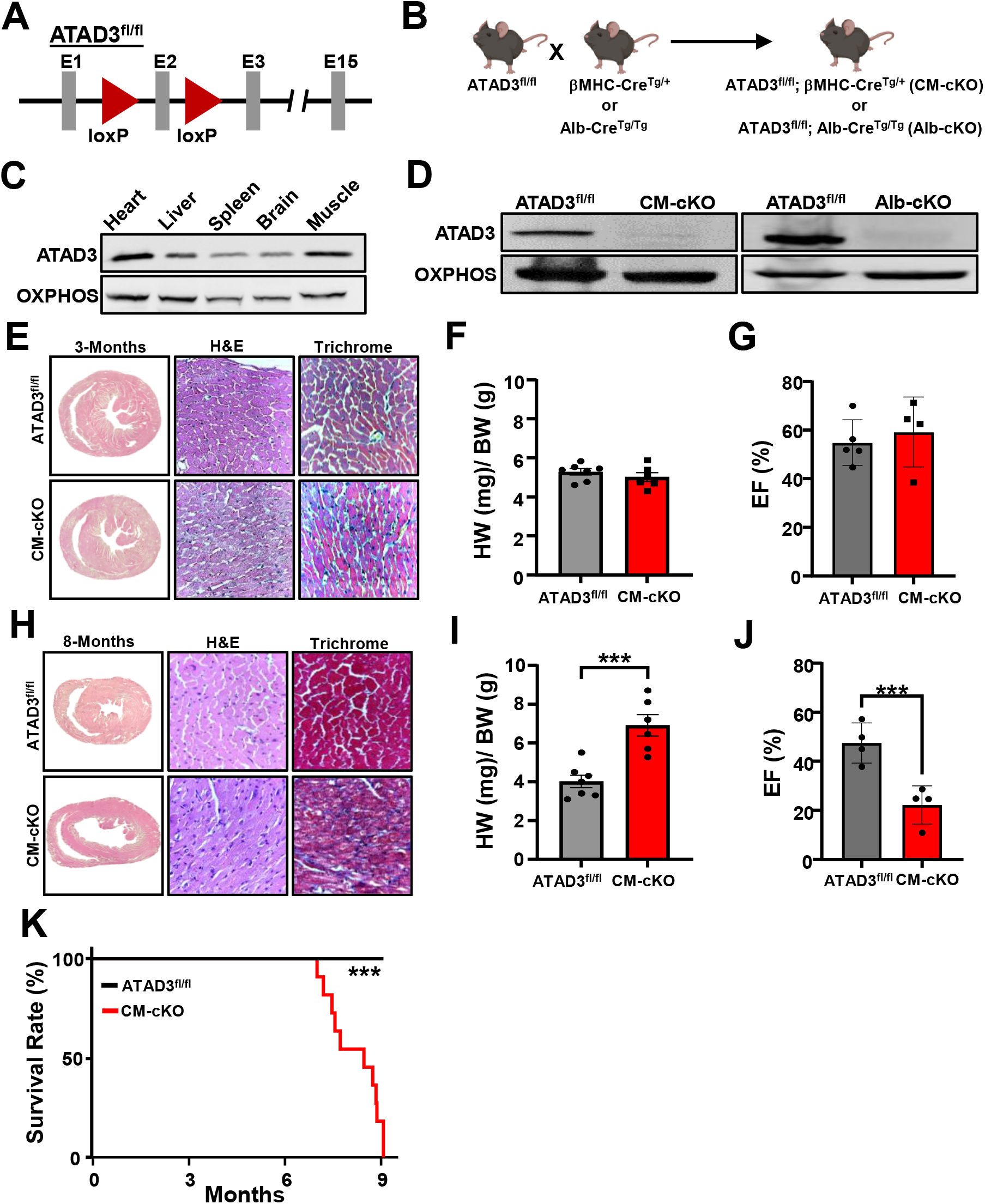
Deletion of Atad3 from the heart results in cardiomyopathy and premature death. (**A**) Schematic representation of the Atad3 gene locus and location of loxP sites. (**B**) Breeding scheme of Atad3^fl/fl^ mice with either βMHC-Cre or Alb-Cre mice transgenic mice to generate cardiomyocyte specific deletion of Atad3 (CM-cKO) or hepatocyte specific deletion of Atad3 (Alb-cKO), respectively. (**C**) Western Blot of ATAD3 using isolated mitochondrial lysates from various labeled tissues from WT mice. (**D**) Western blots for ATAD3 and controls using isolated mitochondrial lysates from heart or livers to confirm genetic deletion. (**E**) Representative gross and histological images of H&E and Masson’s trichrome staining on cross sections of ATAD3^fl/fl^ and CM-cKO hearts at 3-months. (**F**) Quantifications of heart weight over body weight (HW/BW) of ATAD3^fl/fl^ and CM-cKO hearts at 3-months. (**G**) Ejection fraction percentage (EF) % of either ATAD3^fl/fl^ and CM-cKO mice at 3 months. (**H**) Representative gross and histological images of H&E and Masson’s trichrome staining on cross sections of ATAD3^fl/fl^ and CM-cKO hearts at 8 months. (**I**) Quantifications of HW/BW of ATAD3^fl/fl^ and CM-cKO hearts at 8 months. (**J**) EF % of ATAD3^fl/fl^ and CM-cKO mice at 7-8 months. (**K**) Survival curve analysis of ATAD3^fl/fl^ and CM-cKO mice (median survival 8.5 months p<.001 Log-rank (Mantel-Cox test). ***P < 0.001;

To analyze mitochondrial morphology and function, we subjected isolated mitochondria from 3-month-old-mice to TEM imaging and found that ATAD3 null mitochondria are electron dense, but display increased cristae luminal width (Fig. 2A). Additionally, we subjected isolated cardiac mitochondria to a mitochondrial stress test. Here, we found that cardiac mitochondria lacking ATAD3 have a small increase in baseline respiration, but a blunted response to ADP compared to control, however, maximal respiratory capacity remained intact (Fig. 2B). Additionally, we assessed the expression of known mPTP regulating proteins in ATAD3 null cardiac heart mitochondria and determined that every mPTP regulator was unchanged except for a 6-fold increase in BAX and a <2-fold decrease in BAK1 expression (Fig. 2C-H).

**Fig. 2.**
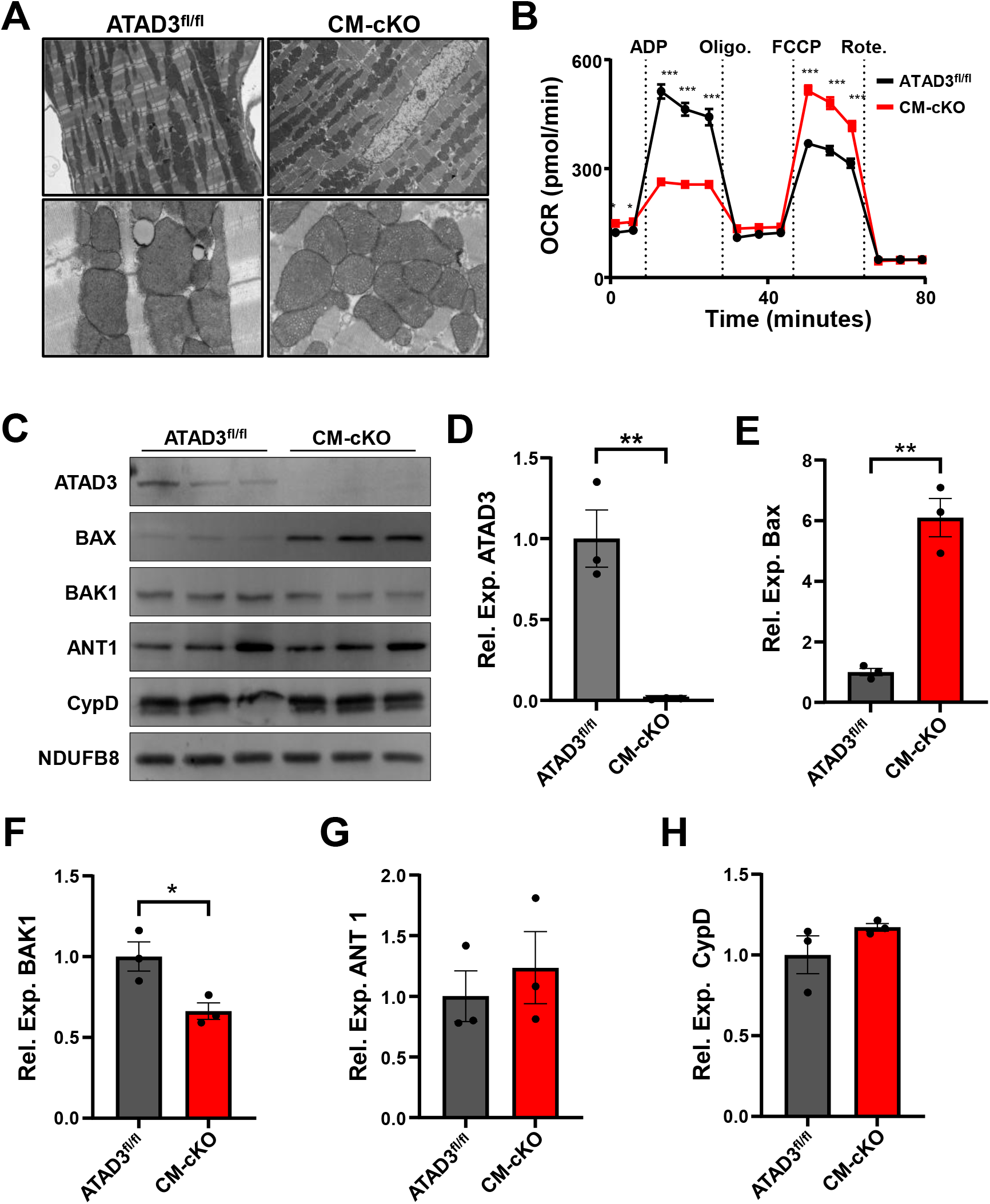
Characterization of cardiac ATAD3 null mitochondria. **(A)** Representative TEM images of mitochondria isolated from either ATAD3^fl/fl^ or CM-cKO hearts. (**B)** Oxygen consumption rate (OCR) of mitochondria isolated from either ATAD3^fl/fl^ or CM-cKO hearts subjected to a mitochondrial stress test. **(C)** Western blots and **(D-H)** quantification of ATAD3, mPTP regulators ANT1 and CypD, and Bcl-2 family members BAX and BAK1. (n=3; all standardized to NDUFB8). *P < 0.05; **P < 0.01; ***P < 0.001.

To determine if ATAD3 plays a role in mPTP opening, we isolated both liver and heart mitochondria from the ATAD3 deleted tissues and controls and subjected them to the mitochondrial Ca^2+^ retention capacity (CRC) and swelling assays. Mitochondria lacking ATAD3 have significantly increased CRC and do not undergo Ca^2+^-induced swelling (Fig 3A-F). Additionally, we visualized these mitochondria using TEM, Ca^2+^ treated control mitochondria undergo cristae dissipation and swelling indicative of mPTP opening, while ATAD3 null mitochondria remain intact and electron dense even when treated with extremely high levels of Ca^2+^ (Fig. 3G). To investigate if these findings were specific to Ca^2+^, we subjected the mitochondria to lipid peroxidation-dependent swelling by tBHP treatment, which is not dependent mPTP opening^18^. ATAD3 null mitochondria swell in response to tBHP treatment similarly to control mitochondria, suggesting specificity for mPTP-dependent swelling (Fig. 2H and I). Cyclosporine A (CsA), through CypD inhibition, and ADP, through the ANT family, are commonly used to desensitize the mPTP to Ca^2+ 19^. Since mPTP opening is inhibited in the absence of ATAD3, we wanted to determine if CsA or ADP would have any effect on ATAD3 null mitochondria. As expected, CsA and ADP significantly increased CRC and delayed mitochondrial swelling in ATAD3 expressing liver and heart mitochondria (Fig. 4A-C and H-J), however, CsA and ADP failed to increase CRC or alter swelling in ATAD3 null liver and heart mitochondria (Fig. 4D-F and K-M).

**Fig. 3.**
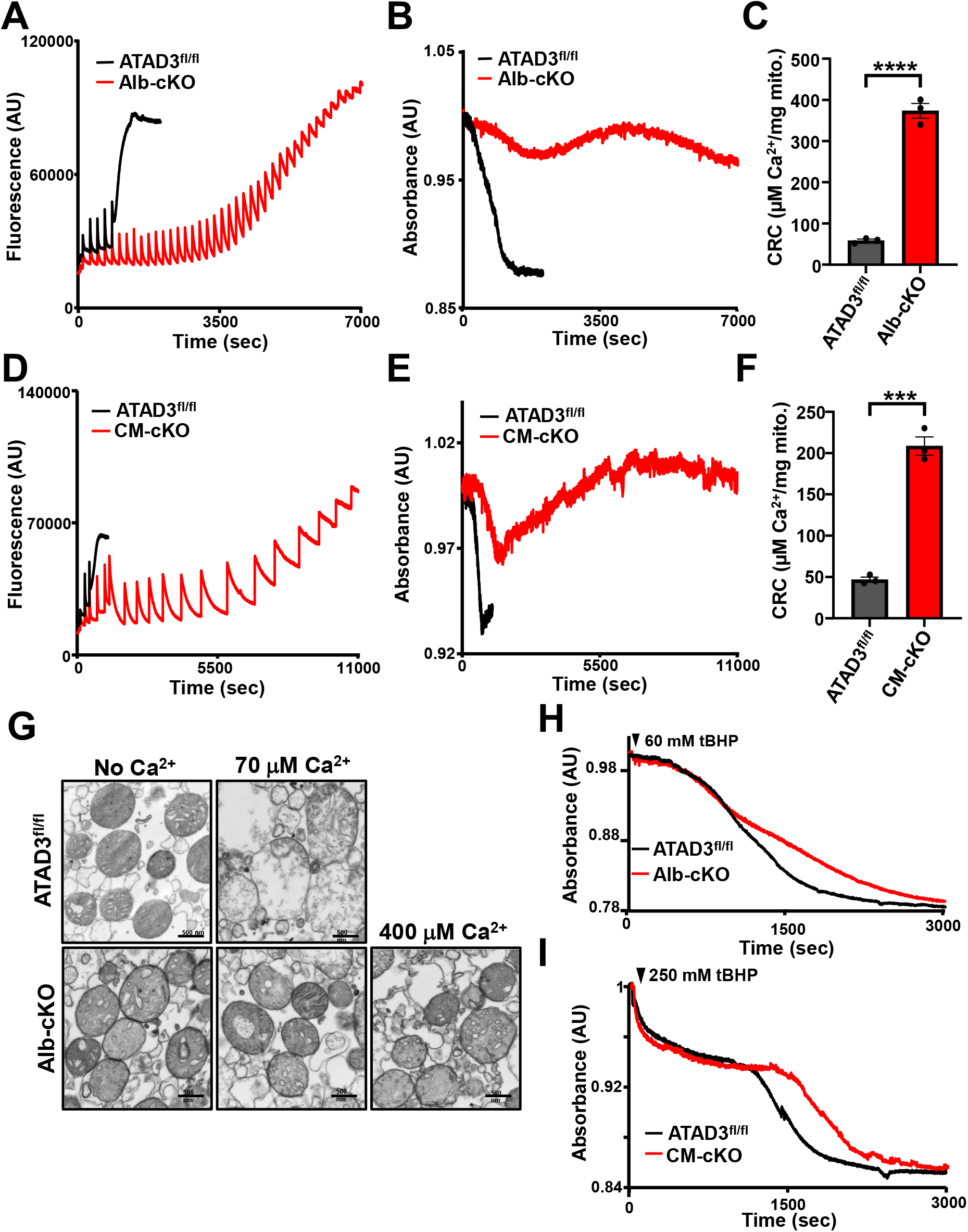
ATAD3 null mitochondria have increased CRC and do not undergo mPTP-dependent swelling. **(A)** Representative traces of either ATAD3^fl/fl^ or Alb-cKO isolated liver mitochondria treated with boluses of calcium (20 µM CaCl_2_). **(B)** Representative trace of mitochondrial swelling corresponding to **(A). (C)** Quantification of CRC calculated from **(A). (D)** Representative traces of either ATAD3^fl/fl^ or CM-cKO isolated cardiac mitochondria treated with boluses of calcium (10µM CaCl_2_). **(E)** Representative trace of mitochondrial swelling corresponding to **(D). (F)** Quantification of CRC calculated from **(D). (G)** Representative traces of either ATAD3^fl/fl^ or Alb-cKO isolated liver mitochondria treated with either no calcium (left) 70 µM calcium (middle) or 400 µM calcium (right). **(H)** Representative swelling traces of either ATAD3^fl/fl^ or Alb-cKO isolated liver mitochondria treated with 60mM tBHP. **(I)** Representative swelling traces of either ATAD3^fl/fl^ or CM-cKO isolated cardiac mitochondria treated with 250mM tBHP. ***P < 0.001; ****P < 0.0001.

**Fig. 4.**
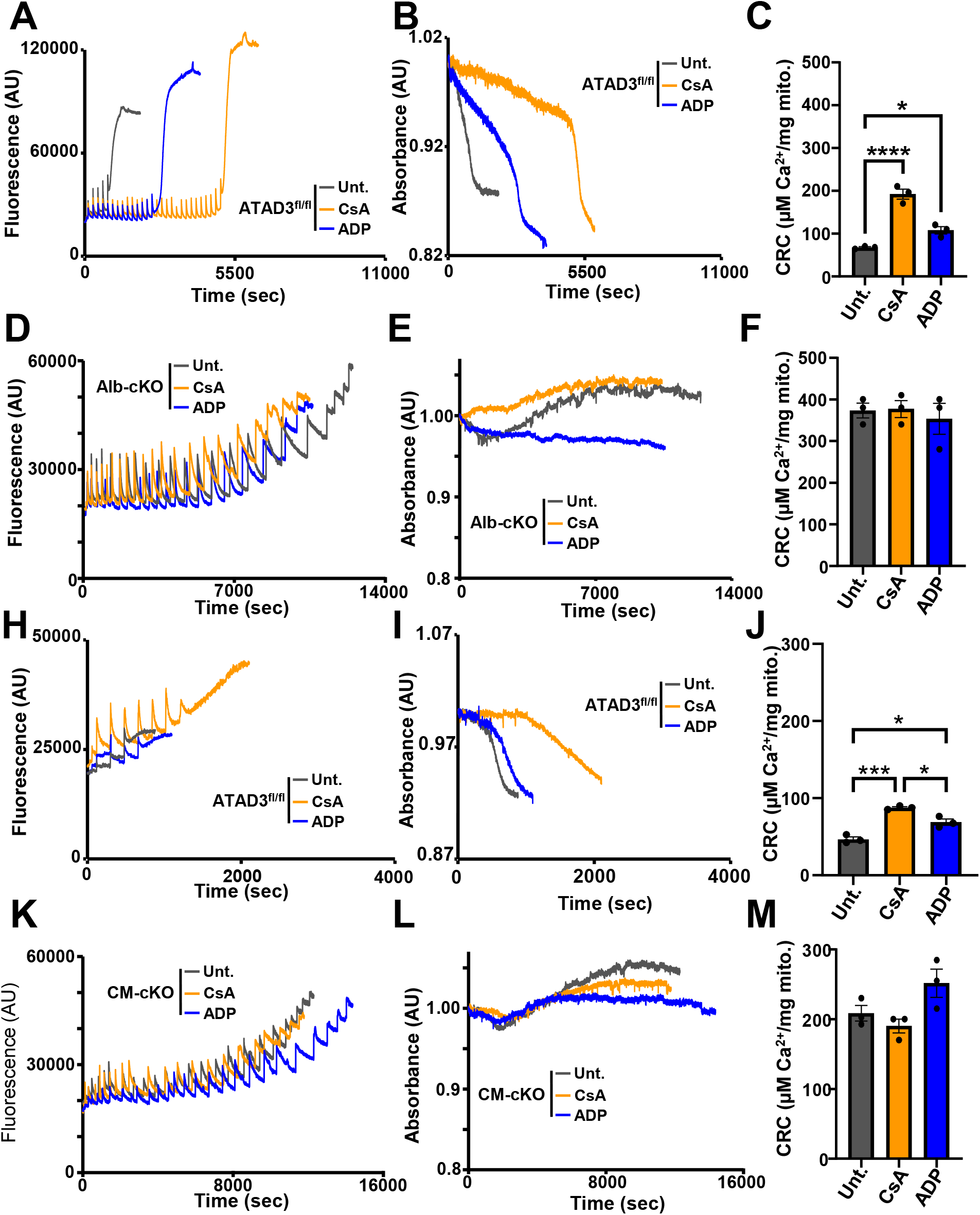
mPTP desensitization by CsA or ADP does not increase CRC in ATAD3 null mitochondria. **(A)** Representative traces of ATAD3^fl/fl^ isolated liver mitochondria pretreated with either 2 µM CsA (blue trace) or 300 µM ADP (orange trace) and treated with boluses of calcium (20 µM CaCl_2_). **(B)** Representative trace of mitochondrial swelling corresponding to **(A). (C)** Quantification of CRC calculated from **(A). (D)** Representative traces of Alb-cKO isolated liver mitochondria pretreated with either 2 µM CsA (blue trace) or 300µM ADP (orange trace) and treated with boluses of calcium (20 µM CaCl_2_). **(E)** Representative trace of mitochondrial swelling corresponding to **(D). (F)** Quantification of CRC calculated from **(D). (H)** Representative traces of ATAD3^fl/fl^ isolated cardiac mitochondria pretreated with either 2 µM CsA (blue trace) or 300µM ADP (orange trace) and treated with boluses of calcium (20 µM CaCl_2_). **I** Representative trace of mitochondrial swelling corresponding to **(H). (J)** Quantification of CRC calculated from **(H)** (n=3). (**K)** Representative traces of CM-cKO isolated cardiac mitochondria pretreated with either 2 µM CsA (blue trace) or 300 µM ADP (orange trace) and treated with boluses of calcium (20 µM CaCl_2_). **(L)** Representative trace of mitochondrial swelling corresponding to **(K). (M)** Quantification of CRC calculated from **(K)**. *P < 0.05; ***P < 0.001; ****P < 0.0001.

Since mPTP activity is abolished upon ATAD3 deletion, we investigated whether ATAD3A possesses intrinsic channel-forming activity. To test this, we reconstituted recombinant human ATAD3A into artificial liposomes and performed patch-clamp recordings. Liposomes containing ATAD3A exhibited robust channel activity, indicating that ATAD3A is capable of forming pores (Fig. 5A–C). The observed conductance changes were consistent with previously reported mPTP properties. Importantly, channel activity was inhibited by ATP, suggesting that ATAD3A-mediated pore formation is ATP-sensitive (Fig. 5A–C). These findings support the hypothesis that ATAD3A is the pore-forming component of the mPTP.

**Fig. 5.**
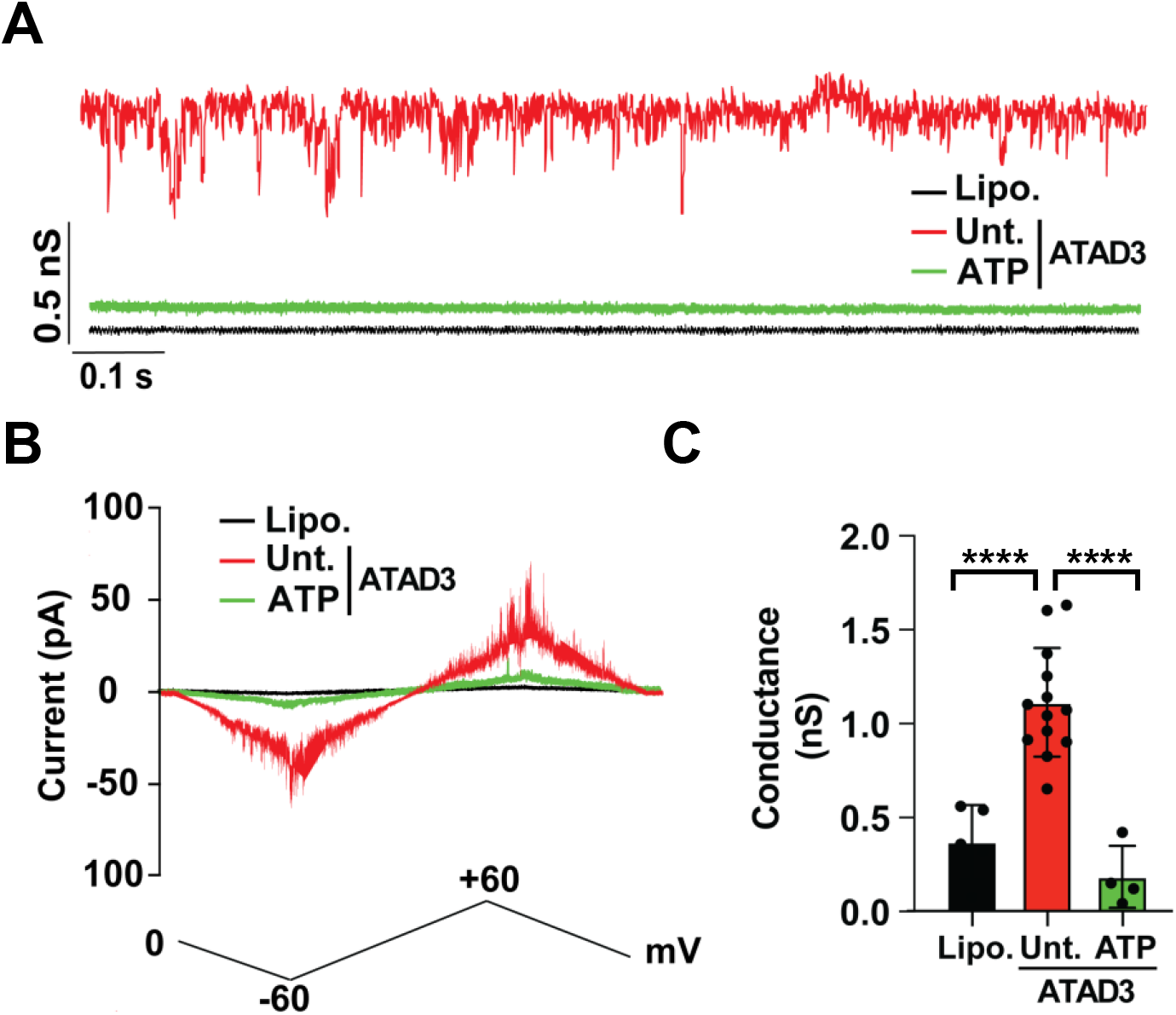
Recombinant ATAD3A exhibits intrinsic channel activity in liposomes that is inhibited by ATP. **(A)** Representative patch-clamp current recordings from liposomes alone (Lipo., black), liposomes reconstituted with untreated recombinant ATAD3a (Unt., red), and ATAD3a in the presence of ATP (green). **(B)** Current-voltage (I-V) relationship showing robust, voltage-dependent conductance in liposomes containing untreated ATAD3a compared to minimal activity in ATP-treated and control liposomes. **(C)** Quantification of single-channel conductance across conditions. ****P < 0.0001.

Inhibition of mPTP-dependent necrosis by CsA protects against I/R injury in mice^4,5^. Here, we investigated if ATAD3 null hearts are protected from I/R injury. We subjected Atad3 CM-cKO and Atad3^fl/fl^ mice to I/R injury (1 hour ischemia followed by 24 hours reperfusion) and then measured infarct size and area at risk. Indeed, deletion of Atad3 from cardiomyocytes significantly reduced infarct size following I/R injury (Fig 6A-C). Furthermore, to demonstrate that Atad3 deletion acts through mPTP inhibition, we pretreated mice with CsA. As expected, CsA treatment on the control Atad3^fl/fl^ animals significantly reduced infarct size compared to the vehicle control animals (Fig 7A-C). However, CsA conferred no additional protection in Atad3 CM-cKO mice, indicating that mPTP-dependent necrosis is already fully suppressed in the absence of ATAD3 (Fig. 7A–C). These findings suggest that ATAD3 deletion protects the heart by inhibiting mPTP opening during I/R injury and that the remaining infarct is mPTP-independent.

**Fig. 6.**
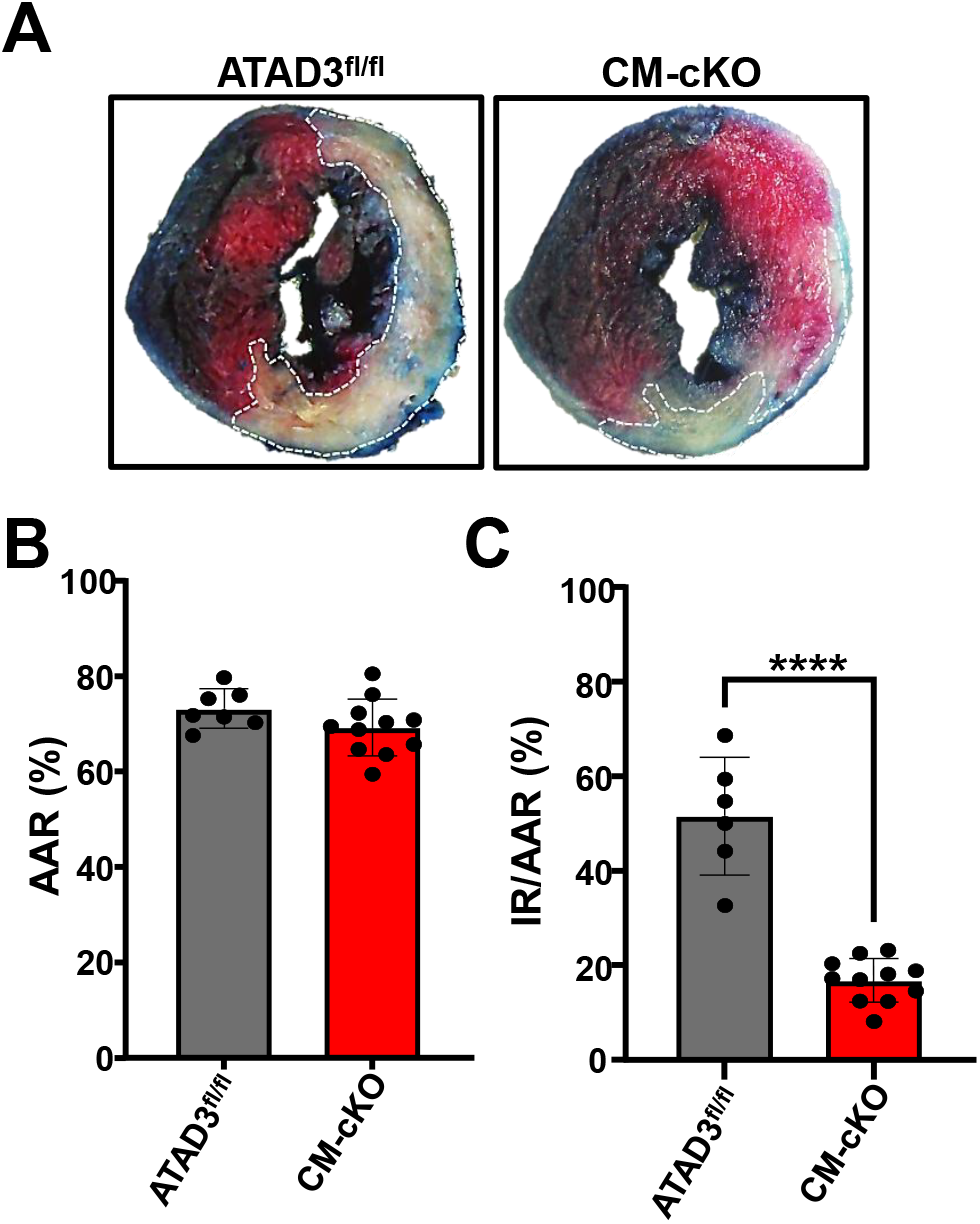
Deletion of Atad3 from the heart protects against I/R injury. **(A)** Representative images of either ATAD3^fl/fl^ or CM-cKO hearts following 1 hour of ischemia and 24 hours reperfusion (blue; Evan’s blue dye: unaffected tissue, TCC staining, red (healthy tissue) and white (infarct) **(B)** Quantification of area at risk (AAR) percentage. **(C)** Quantification of infarct region (IR) over AAR. ****P < 0.0001

**Fig. 7.**
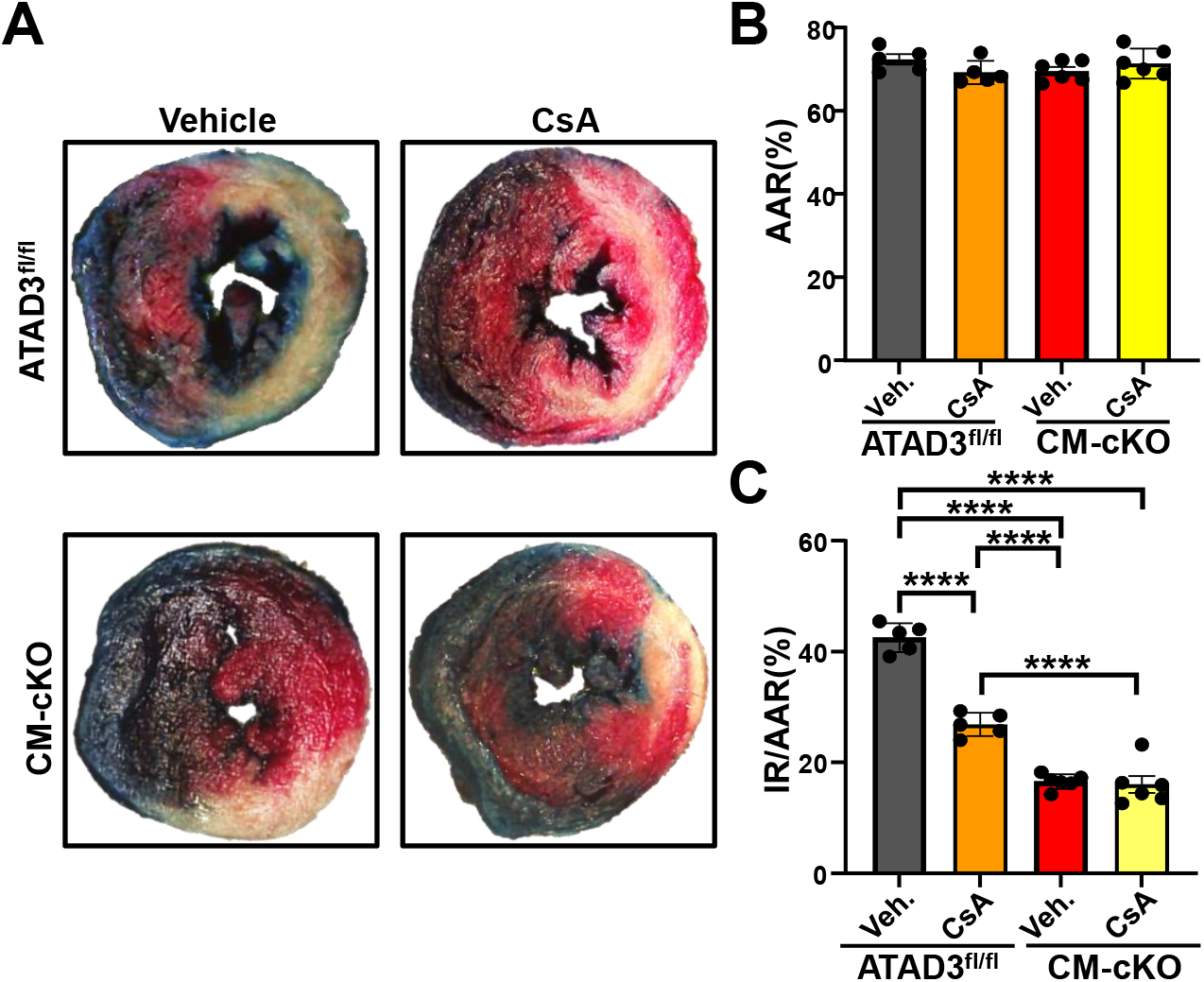
CsA does not provide further protection against I/R injury in Atad3 null hearts. **A** Representative images of either Evan’s blue dye and TTC stained ATAD3^fl/fl^ or CM-cKO hearts treated with either vehicle treated (left) or 10 mg/kg CsA treated (right) followed by 1 hour of ischemia and 24 hours. **(B)** Quantification of the area at risk (AAR) percentage. **(C)** Quantification of infarct region (IR) over AAR. ****P ≤ 0.0001

## Discussion

The molecular identity of the mitochondrial permeability transition pore (mPTP) has remained elusive since its discovery. To date, only non-essential regulators of the mPTP, such as the ANT family, Bax and Bak1, and CypD, have been characterized^3^. Here, we demonstrate that ATAD3 is an essential component of the mPTP. Deletion of ATAD3 blocks Ca^2+^-induced mitochondrial swelling, enhances mitochondrial CRC, and confers protection against I/R injury. To our knowledge, this is the first description of an essential component of the mPTP.

Beyond the data presented here, the structural and functional characteristics of ATAD3 make it a compelling candidate for an essential component of the mPTP. ATAD3 is an integral inner mitochondrial membrane (IMM) protein that spans the intermembrane space^12^. Early descriptions of the mPTP proposed it as a continuous structure bridging the IMM and outer mitochondrial membrane (OMM), and subsequent studies have localized the pore to contact sites between these membranes^20^. It is well established that OMM proteins, including Bax, Bak1, and other Bcl-2 family members, modulate the sensitivity of the inner membrane component of the mPTP to matrix Ca^2+10,21^. Thus, ATAD3 may function as a scaffold that links OMM regulators to IMM components of the pore. This is supported by our observation that ATAD3 deletion results in increased expression of Bax, suggesting a compensatory response. Additionally, ATAD3 forms oligomers via its coiled-coil domains, and such oligomerization has been implicated in Alzheimer’s disease, where its inhibition reduces disease burden^17,22^. It is plausible that ATAD3 oligomerization enables pore formation within the IMM, making it a candidate for the pore-forming unit of the mPTP. Finally, ATAD3 contains a large matrix-facing domain rich in proline residues (nine in total), which may be relevant given that CypD facilitates mPTP opening through its prolyl isomerase activity.

The physiological relevance of ATAD3 is further underscored by human genetics. Humans express three ATAD3 isoforms (ATAD3A, ATAD3B, and ATAD3C) with ATAD3A being the most abundant and functionally critical in most tissues^12^. Mutations in the ATAD3A locus are associated with a spectrum of mitochondrial diseases, most notably Harel-Yoon syndrome, a neurodevelopmental disorder characterized by global developmental delay, hypotonia, cerebellar atrophy, and progressive neurological decline^23^. These clinical phenotypes reflect profound mitochondrial dysfunction and highlight the essential role of ATAD3A in maintaining mitochondrial structure and function. Importantly, many of the pathogenic variants identified in patients cluster within domains critical for ATAD3A oligomerization or ATPase activity, suggesting that disruption of these features compromises its ability to regulate mitochondrial dynamics, nucleoids, and potentially, mPTP activity^24^. Given our data showing that loss of ATAD3 abolishes mPTP opening, these disease phenotypes may in part be driven by impaired mPTP function, leading to defective Ca^2+^ handling, resistance to regulated necrosis, or aberrant mitochondrial signaling. Thus, ATAD3A is not only essential for basal mitochondrial homeostasis, but may also be a key player in cell death pathways across a range of physiological and pathological contexts. These findings broaden the functional landscape of ATAD3 and suggest that dysregulation of mPTP activity may underlie, or contribute to, the pathogenesis of ATAD3A-related mitochondrial diseases.

Taken together, these structural, functional, and genetic data support a model in which ATAD3 is not only essential for mitochondrial organization but may also form the core of the mPTP. This positions ATAD3 as a promising focal point for future mechanistic studies and potential therapeutic targeting in diseases involving mPTP-dependent necrosis.

## Methods

### Animal Models

Atad3 conditional ready/knockout embryonic stem cells were obtained through the Knockout Mouse Project (KOMP) repository. Once these mice were generated, heterozygote Atad3 null mice were crossed to a flp recombinase transgenic mouse to generate the Atad3 floxed allele. To generate cardiac-specific deletion of ATAD3^fl/fl^ mice, we crossed in mice containing the Cre recombinase under the control of βMHC (β-myosin heavy chain) promoter. To generate liver-specific deletion of ATAD3^fl/fl^ mice, we crossed in mice containing the Cre recombinase under the control of Alb (albumin) promoter. All experimental procedures with animals were approved by the Institutional Animal Care and Use Committee (IACUC) of Baylor College of Medicine (protocol IACUC AN-7915). All mice were treated humanely as per compliance with the rules and regulations of animal care and euthanasia under this committee. The minimal number of mice was used in this study to attain statistical significance based on power analysis calculations.

### Mitochondrial isolations

Heart ventricles or livers were isolated from ATAD3^fl/fl^ and CM-cKO mice and washed in a mitochondrial isolation buffer (225 mM mannitol, 75 mM sucrose, 5 mM Hepes, and 1 mM EGTA). Hearts were then minced into 1-2 mm pieces in 1.5 mL mitochondrial isolation buffer. The tissue was homogenized using a Teflon/glass tissue grinder. All steps were performed on ice. The homogenates were then centrifuged at 800g for 5 minutes. The supernatants were collected and centrifuged at 7,500g for 10 minutes. The supernatants were aspirated and the remaining pellets were re-suspended and washed with 7 mL of the mitochondrial isolation buffer. The re-suspended solution was centrifuged at 7,500g for 10 minutes. All centrifugations were performed at 4°C. The supernatant from the final spin and the remaining pellet was re-suspended in 500 µL of a KCl buffer (125 mM KCl, 20 mM Hepes, 2 mM KH_2_PO_4_ and 40µM EGTA (pH 7.2). Mitochondrial concentration was measured using Nanodrop at 280nm absorbance.

### CRC and mitochondrial swelling assays

Two milligrams of isolated mitochondria were suspended in a total volume of 1 ml consisting of a KCl buffer, 1 mM malic acid (Sigma-Aldrich), 7 mM pyruvate (Sigma-Aldrich), and 50 nM Calcium Green-5N (Invitrogen) in a quartz cuvette, which was placed inside the fluorimeter (PTI QuantaMaster 800, Horiba Scientific). Calcium uptake was measured by fluorescence emission of Calcium Green-5N. Simultaneously, mitochondrial swelling was measured by transmitted light.

Some experiments included ADP (300 μM) (Sigma-Aldrich, A2754) or CsA (2 μM) (Sigma-Aldrich, 30024) to desensitize the mPTP. CaCl_2_ (20 μM)(Sigma-Aldrich, C4901) was added into this system in succession until mPTP opening occurred, indicated by an increase in Calcium Green-5N fluorescence or when mitochondria were saturated with Ca^2+^ and were no longer able to take up further additions of CaCl_2_, indicated by a stair stacking of Calcium Green-5N fluorescence. For tBHP swelling analysis either 60 mM (for livers) or 250 mM of tert-Butyl hydroperoxide (tBHP) (Sigma-Aldrich; 458139) was added to the cuvette.

### Transmission Electron Microscopy

For transmission electron microscopy of heart mitochondria, hearts from ATAD3^fl/fl^ and CM-cKO were isolated and washed in Hank’s Balanced Salt Solution. Hearts were then minced into 1-2 mm pieces before being incubated in EM grade 4% paraformaldehyde (Electron Microscopy Sciences, 15713-S) overnight at 4° C. Microscope imaging was performed by the Electron Microscopy Core Facility at UT Southwestern Medical Center. For TEM of liver mitochondria, mitochondria from ATAD3^fl/fl^ and Alb-cKO mice were isolated and 2 mg of mitochondria were added to a 1.5 mL Eppendorf tube to a total volume of 1 mL in KCl buffer. These samples were treated with boluses of calcium in parallel to the CRC assay until mPTP opening was observed. Additionally, mitochondria from Alb-cKO mice were treated with boluses of calcium in parallel to CRC assay until stair stacking was observed. All samples were centrifuged at 5000g for 10 minutes at 4°C. Supernatant was aspirated and EM grade 4% paraformaldehyde was added carefully to not disturb the pellet.

### Western Blotting

Mitochondrial pellets were suspended in a radioimmunoprecipitation assay buffer [10 mM tris-HCl (pH 7.49), 100 nM NaCl, 1 mM EDTA, 1 nM EGTA, 1% Triton X-100, 10% glycerol, 0.1% SDS, and 0.5% sodium deoxycholate] containing protease inhibitor cocktails (Roche). Then, these samples were sonicated before centrifugation (21,000g for 10 min at 4°C). Afterward, the supernatant fractions were diluted in an SDS sample buffer [250 mM tris-HCl (pH 7.0), 10% SDS, 5% β-mercaptoethanol, 0.02% bromophenol blue, and 30% glycerol] before boiling at 100°C for 5 min. Protein samples were then loaded onto 12 to 15% acrylamide gels and then transferred onto polyvinylidene fluoride transfer membranes (MilliporeSigma). The following primary antibodies were used: ANT1 (Signalway, 32484 1:500), ATAD3 (Abnova, H00055210-D01P 1:1000), Bak (EMD Millipore, 06-536 1:800), Bax (Cell Signaling Technology, 2772S 1:1000), CypD (Abcam, ab110324 1:1000), and Total OXPHOS rodent WB antibody cocktail (Abcam, ab110413 1:10,000). These blots were incubated in their respective secondary antibodies: ATAD3, Bax, ANT1, Bak; goat anti-rabbit immunoglobulin G (H+L) (Novus, NB7157 1:15,000); CypD and OXPHOS; goat anti-mouse immunoglobulin G (H+L) (Novus, 7536) for 2 hours at room temperature before being washed with 1xTBST for 5 washes at 6 minutes each. These blots were then incubated in an ECF substrate for 1 minute each and exposed using the Thermo Fisher Scientific iBright imaging system.

### Mitochondrial Stress Test

Mitochondria from ATAD3^fl/fl^ and CM-cKO mouse hearts were isolated in mitochondria isolation buffer (210 mM d-mannitol, 70 mM sucrose, 5 mM Hepes (pH 7.2), 1 mM EGTA, and 0.1% bovine serum albumin (BSA)), and the resulting mitochondrial pellet was re-suspended in 500 μl of a mitochondrial assay buffer (220 mM d-mannitol, 70 mM sucrose, 10 mM KH_2_PO_4_, 5 mM MgCl_2_, 2 mM Hepes (pH 7.2), 1 mM EGTA, and 0.02% BSA); mitochondrial concentration was measured using Bradford assay. OCR was analyzed using the Seahorse XF Extracellular Flux Analyzer (Agilent Technologies). Four micrograms of mitochondria per well were plated on XF96 microplates (Agilent Technologies). The mitochondria were then brought up in an isolation buffer with 0.02% BSA and 5 mM pyruvate and 0.5 mM malic acid. Basal respiration was measured before treatments with 500 μM ADP, 10 μM oligomycin, 5 μM carbonyl cyanide p-trifluoromethoxyphenylhydrazone, and then 1 μM rotenone (all diluted in an isolation buffer with 5 mM pyruvate and 0.5 mM malic acid).

### I/R surgery

Mice were placed in an anesthesia induction chamber followed by a nosecone with isoflurane flowing at 5% at 1 to 2% oxygen and locally anesthetized using bupivacaine (1 to 2 mg/kg) prior to a 1-cm ventral incision on the neck to intubate the trachea with more global anesthetic (isoflurane), following a left lateral thoracotomy (1-cm incision) for isolation of the anterolateral heart. Next, the LAD artery was ligated for 60 min via a slipknot followed by 24-hour reperfusion. Mice were then euthanized, and hearts were excised; the sutures were retied around the LAD artery, and hearts were injected and stained with Evans blue dye and then sliced into 1-mm-thick cross sections and incubated and stained with 2,3,5-triphenyltetrazolium to visualize and quantify the area at risk and infarct regions. Before surgery, some mice were intraperitoneally injected with CsA (10 mg/kg; 2.5% CsA (10 mg/kg) in dimethyl sulfoxide (DMSO), 29.5% polyethylene glycol, molecular weight 300 (PEG-300), 5% Tween 80, and 63% saline) or vehicle (2.5% DMSO, 29.5% PEG-300, 5% Tween 80, and 62% saline) for four consecutive days. On the fourth day of injections, mice were subjected to the surgery.

### Echocardiography

Prior to analysis, the chests of ATAD3^fl/fl^ and CM-cKO were shaved. Measurements were taken on anaesthetized mice (1.5% isoflurane) using VisualSonics F2 Ultrasound 15-MHz microprobe. Both M-mode and B-mode echocardiography were performed. Quantification of ejection fraction (EJ) were performed using Vevo LAB program for M-mode in triplicate per mouse and averaged.

### Histological Analysis

Hearts of either ATAD3^fl/fl^ and CM-cKO were isolated and incubated in 10% neutral buffered formalin (Sigma-Aldrich HT501128) overnight at room temperature. Samples were then dehydrated in 70% ethanol. Samples were then submitted to the Human Tissue Acquisition and Pathology core at Baylor College of Medicine for subsequent paraffin embedding and hematoxylin and eosin (H&E), Masson’s trichrome, and TUNEL staining. Slides were imaged using the Echo Revolve microscope.

### Statistical Analysis

The data presented represent the mean with the error bars representing the SEM. When comparing two groups, and unpaired two-tailed Student’s t test was performed. When comparing multiple groups to identical controls, a one-way analysis of variance (ANOVA) was performed followed by Dunnett’s correction of multiple t testing. All values were considered to be statistically significant when p<.05 as labeled on figure legends. Each biological replicate is represented as a single dot within histograms.

## Author contributions

Conceptualization: JK

Methodology: JDM, PMP, JK

Investigation: DR, SJL, AM, DR, SL, PMP, JK

Visualization: DR, JK

Funding acquisition: JK

Supervision: JK

Writing – original draft: JK

Writing – review & editing: DR, JDM, PMP, JK

### Data and materials availability

All data are available in the main text or the supplementary materials, the ATAD3a^fl/fl^ mice are available upon request.

